# Developmental stage-specific distribution of macrophages in mouse mammary gland

**DOI:** 10.1101/746511

**Authors:** Teneale A. Stewart, Katherine Hughes, David A. Hume, Felicity M. Davis

## Abstract

Mammary gland development begins in the embryo and continues throughout the reproductive life of female mammals. Tissue macrophages (Mϕs), dependent on signals from the Mϕ colony stimulating factor 1 receptor (CSF1R), have been shown to regulate the generation, regression and regeneration of this organ, which is central for mammalian offspring survival. However, the distribution of Mϕs in the pre- and post-natal mammary gland, as it undergoes distinct phases of development and regression, is unknown or has been inferred from immunostaining of thin tissue sections. Here, we used optical tissue clearing and 3-dimensional imaging of mammary tissue obtained from *Csf1r-EGFP* mice. Whilst tissue Mϕs were observed at all developmental phases, their abundance, morphology, localization and association with luminal and basal epithelial cells exhibited stage-specific differences. Furthermore, sexual dimorphism was observed at E14.5, when the male mammary bud is severed from the overlying epidermis. These findings provide new insights into the localization and possible functions of heterogeneous tissue Mϕ populations in mammogenesis.

## 1 Introduction

Mammary gland development is phasic, with distinct developmental periods occurring in the embryo, at puberty and during pregnancy/lactation (Lloyd-Lewis et al., 2017; Watson and Khaled, 2008). The formation of the milk lines occurs at approximately embryonic day (E) 10 in mice and within 36 h resolves into five pairs of disk-shaped thickenings known as mammary placodes (Cowin and Wysolmerski, 2010). At around E12.5, mammary placodes invaginate into the dermal mesenchyme forming the mammary buds, which later elongate and invade the fat pad precursor, creating a rudimentary epithelial tree (Cowin and Wysolmerski, 2010; Lilja et al., 2018; Paine and Lewis, 2017). During embryonic development, multi-potent mammary stem cells are replaced by unipotent luminal and basal stem/progenitor cells (Lilja et al., 2018; Wuidart et al., 2018), with epithelial cell identities being resolved by E15.5 (Lilja et al., 2018).

Initial postnatal growth of the mammary epithelium is proportional to body size and it is not until puberty that ductal elongation occurs, fueled by proliferation of adult mammary stem/progenitor cells within terminal end bud (TEB) structures (Davis et al., 2016; Lloyd-Lewis et al., 2017, 2018; Paine and Lewis, 2017). Further epithelial expansion occurs during pregnancy to generate the functional (milk-producing) alveolar epithelium (Davis et al., 2016; Watson and Khaled, 2008). With the cessation of infant suckling, alveolar mammary epithelial cells undergo massive programmed cell death (a process known as post-lactational involution), returning the mammary gland to a near pre-pregnant state that is capable of supporting future pregnancies (Lloyd-Lewis et al., 2017; Sargeant et al., 2014).

Mϕs are present in all adult tissues (Hume et al., 2019b). These cells are first and foremost professional phagocytes, but also regulate tissue development, function and dysfunction (Hume, 2015; Naik et al., 2018; Yang et al., 2018). In the normal postnatal mammary gland, Mϕs regulate ductal morphogenesis during puberty (Gouon-Evans et al., 2000; Ingman et al., 2006; Van Nguyen and Pollard, 2002), alveolar budding during ovarian cycling (Chua et al., 2010), alveologenesis in pregnancy (Pollard and Hennighausen, 1994) and tissue remodeling during post-lactational involution (Hughes et al., 2012; O’Brien et al., 2010, 2012), with many of these processes being impaired in mice that lack colony stimulating factor 1 (CSF1). Moreover, Mϕs identified by fluorescence-activated cell sorting (FACS) of disaggregated tissue were detected within the embryonic mammary gland by E16.5 and fetal-derived Mϕs were apparently retained and expanded by self-renewal in adult mammary tissue (Jäppinen et al., 2019).

With accumulating evidence demonstrating the dependence of the mammary epithelium on Mϕs at all developmental stages, it is tempting to speculate that tissue-resident Mϕs institute or influence a putative mammary stem cell niche, as has been shown for hematopoietic stem cells (Winkler et al., 2010), intestinal stem cells (Sehgal et al., 2018) and hair follicle stem cells (Castellana et al., 2014; Naik et al., 2018). Indeed, the activity of mammary “stem” or repopulating cells (defined as a subset of basal cells that are capable of recreating the bi-layered mammary epithelium upon limiting dilution transplantation) is reduced when cells are transplanted into the cleared fat pads of Mϕ-depleted recipient mice (Gyorki et al., 2009). More recently, mammary repopulating cells were shown to express a Notch ligand Delta like 1 (DLL1) and *Dll1*-conditional knockout mice showed reduced mammary repopulating activity and lower levels of F4/80^+^ Mϕs (Chakrabarti et al., 2018). Thus, it has been suggested that DLL1-expressing basal cells activate Notch-expressing Mϕs in a reciprocal stem cell-macrophage niche (Chakrabarti et al., 2018; Kannan and Eaves, 2018). Studies revealing developmental stage-dependent distribution of Mϕs in the mammary gland, including their sites of confluence, would provide further evidence for the existence of a stem cell-macrophage niche in this organ and may help to reveal the specific and stage-dependent localization of mammary stem/progenitor cells within the dynamic, bilayered epithelium under physiological conditions. Here, we utilize a fluorescent reporter model and optical tissue clearing techniques to reveal the presence, prevalence and position of Mϕs in the mammary gland at all phases of development.

## 2 Materials and Methods

### 2.1 Reagents

Neutral buffered formalin (NBF), Quadrol®, triethanolamine and 4’,6-diamidino-2-phenylindole (DAPI) dilactate were purchased from Sigma Aldrich. Normal goat serum was purchased from ThermoFisher. Urea and sucrose were purchased from Chem-Supply. Triton-X-100 was purchased from VWR International. The following primary antibodies were used for immunostaining: chicken anti-GFP (Abcam, ab13970, batch #s GR3190550-3 and −12), rat anti-F4/80 (Novus, NB600-404), rat anti-keratin 8 (DSHB, TROMA-I, batch #s 7/7/16 and 30/3/17), rabbit anti-keratin 5 (BioLegend, 905504, batch # B230397) and rabbit anti-SMA (Abcam, ab5694, batch # GR3183259-26). The following secondary antibodies were used: goat anti-chicken Alexa Fluor-488 (ThermoFisher, A21236), goat anti-rat Cy3 (ThermoFisher, A10522) and goat anti-rabbit Alexa Fluor-647 (ThermoFisher, A21245).

### 2.2 Animal models

Animal experimentation was carried out in accordance with the *Australian Code for the Care and Use of Animals for Scientific Purposes* and the *Queensland Animal Care and Protection Act (2001)*, with local animal ethics committee approval. Animals were housed in individually-ventilated cages with a 12 h light/dark cycle. Food and water were available *ad libitum*. *Csf1r-EGFP* (MacGreen) (Sasmono et al., 2003) mice were a kind gift from A/Prof Allison Pettit (Mater Research Institute-UQ). Mice were maintained as hemizygotes on a C57BL6/J background. C57BL6/J mice were obtained from the Animal Resources Centre (Western Australia).

To obtain mammary tissue during gestation, female mice were mated and tissue harvested 14.5 days-post-coitus. GFP^+^ embryos (E14.5) were also harvested and analyzed after PCR-sexing. To obtain tissue during lactation, female mice were mated, allowed to litter naturally and lactating mammary tissue harvested on day 10 of lactation. For studies during involution, females were allowed to nurse for 10 days and mammary glands harvested 96 h post forced involution. Mammary glands from pre-pubertal female GFP^+^ mice (postnatal day 10) were also harvested and analyzed.

### 2.3 CUBIC-based tissue clearing and IHC

Tissue clearing was performed as previously optimized and described (Davis et al., 2016; Lloyd-Lewis et al., 2016). Briefly, mammary tissue was spread on foam biopsy pads and fixed for 6-9 h in NBF (10%). Embryos were fixed whole. For CUBIC-based clearing, tissue was immersed in Reagent 1A (Lloyd-Lewis et al., 2016; Susaki et al., 2014) at 37°C for 3 days before washing and blocking in goat serum (10%) in PBS with Triton-X-100 (0.5%) overnight at 4°C. Tissue was incubated in primary antibody in blocking buffer for 4 days and secondary antibody in blocking buffer for 2 days at 4°C. DAPI (5 μg/mL) treatment was performed for 2-3 h at room temperature (omitted for second harmonic generation) and tissue was immersed in modified Reagent 2 (Lloyd-Lewis et al., 2016) at 37°C for at least 24 h prior to imaging.

### 2.4 Immunohistochemistry (FFPE slides)

IHC on FFPE slides was performed as previously described in detail (Stewart et al., 2019). Wholemount immunostaining using anti-GFP antibody was performed prior to processing for paraffin embedding.

### 2.5 Microscopy

Immunostained tissue sections were imaged using an Olympus BX63 upright epifluorescence microscope using UPlanSAPO 10×/0.4. 20×/0.75, 40×/0.95, 60×/1.35 and 100×/1.35 objective lenses. Immunostained optically-cleared tissue was imaged using an Olympus FV3000 laser scanning confocal microscope with UPLSAPO 10×/0.40, UPLSAPO 20×/0.75, UPLSAPO 30×/1.05 and UPLFLN 40×/0.75 objective lenses. 3D de-noising was performed as previously described (Boulanger et al., 2010). For SHG, images were acquired using a Mai Tai DeepSee multiphoton laser on a Zeiss 710 laser scanning inverted microscope. Visualization and image processing was performed in ImageJ (v1.52e, National Institutes of Health) (Linkert et al., 2010; Schindelin et al., 2012).

## 3 Results

### 3.1 Mϕs are present in the embryonic bud and early postnatal gland with sexually dimorphic distribution

Mϕs have never been visualized in the embryonic mammary gland and, until recently, were thought to arrive postnatally (Jäppinen et al., 2019). A study by Jäppinen et al. has revealed the presence of F4/80^+^ cells in digested mammary tissue by E16.5 by flow cytometry (Jäppinen et al., 2019). However, in the absence of *in situ* imaging, it is currently unclear whether these embryonic Mϕs physically associate with the developing mammary epithelium, as has been observed in the postnatal gland.

To assess Mϕ distribution in 3-dimensions in intact mammary tissue, we used a *Csf1r-EGFP* mouse model (Sasmono et al., 2003), combined with methods for optical tissue clearing and deep tissue imaging (**Supplementary Fig. 1**) (Davis et al., 2016; Lloyd-Lewis et al., 2016, 2018) (Lloyd-Lewis et al., manuscript in preparation). In this model, green fluorescent protein (GFP) expression in tissues is restricted to monocytes and Mϕs in the developing embryo, starting with yolk sac-derived phagocytes, and in all adult tissues (Hume et al., 2019a; Sasmono et al., 2003). Much lower expression in granulocytes and some B lymphocytes is detectable by FACS, but not in tissues. Multi-color fluorescence immunostaining of tissue sections from mouse spleen confirmed that the majority of GFP^+^ cells were also positive for the Mϕ cell surface marker, F4/80 (**Supplementary Fig. 2**).

In 3D image stacks of female *Csf1r-EGFP* embryos, Mϕs were detected in the mammary and dermal mesenchyme surrounding the mammary epithelial bud as early as E14.5 (**Fig. 1A** and **Supplementary Fig. 3A**). As expected (Sasmono et al., 2003), Mϕs were also present in the embryonic liver at this stage (**Fig. 1B**), and it has been suggested that these fetal liver-derived Mϕs contribute extensively to the pool of tissue Mϕs present in the adult gland (Jäppinen et al., 2019). Our data show that Mϕs were positioned adjacent to the embryonic mammary epithelium around the time of lineage segregation (Lilja et al., 2018; Wuidart et al., 2018). Interestingly, although Mϕs were positioned around the embryonic bud, they were rarely observed to directly interact with the developing epithelium of female embryos (**Fig. 1A** **and Supplementary Fig. 3A**). In contrast, Mϕs directly contacted and invaded the mammary bud of male mice at E14.5, the developmental period when the male bud is severed from the overlying epidermis in mice and begins to regress (**Fig. 1C-D** and **Supplementary Fig. 3B**) (Cowin and Wysolmerski, 2010; Dunbar et al., 1999; Heuberger et al., 2006). Mammary Mϕs were also observed in the early postnatal period in female mice (**Fig. 1E-F)**. By this stage, however, Mϕs were positioned around and inside of this rudimentary structure, apparently interacting with the epithelium (**Fig. 1E**).

**Fig. 1:**
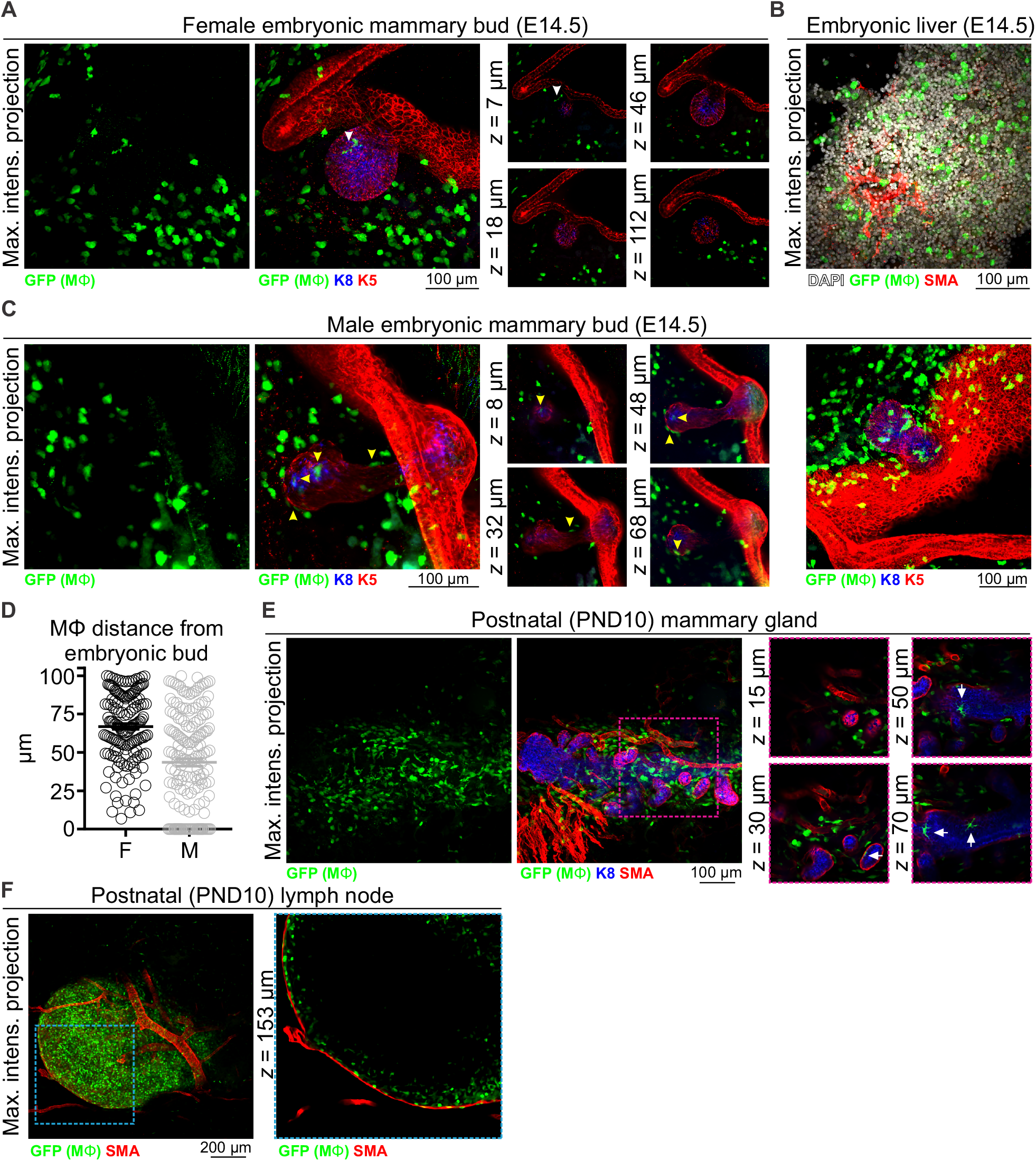
Mϕs in the embryonic and early postnatal mouse mammary gland. Maximum intensity *z*-projection and individual optical slices of cleared tissue from (**A-B**) embryonic (E14.5) female mice and (**C**) embryonic (E14.5) male mice. (**D**) The distance of Mϕs (within a 100 μm radius) of the female and male embryonic buds. Mϕs contacting the bud or inside of the bud were assigned a value of 0; this was only observed in male embryos. (**E**) Mammary tissue from postnatal day (PND) 10 *Csf1r-EGFP* female mice. (**F**) Inguinal lymph node from PND10 mice showing subcapsular sinus Mϕs. Keratin (K) 8 immunostaining shows K8-positive luminal cells; K5 immunostaining reveals K5-expressing basal cells; smooth muscle actin (SMA) immunostaining reveals basal cells and SMA-positive vessels. White arrowhead in (A) points to a Mϕ that appears to be in contact with the embryonic bud in the maximum intensity projection, but is revealed to be positioned in the mammary mesenchyme above the bud in optical slices. Yellow arrowheads in (C) point to Mϕs that are in direct contact with the embryonic bud. Arrows in (E) point to Mϕs that are in contact with the PND10 mammary epithelium. Images are representative of 3 mice/embryos at each developmental stage.

### 3.2 Mϕs envelope and infiltrate the elongating terminal end bud during ductal morphogenesis

Mϕs are essential for normal ductal morphogenesis during puberty (Gouon-Evans et al., 2000; Ingman et al., 2006; Van Nguyen and Pollard, 2002). Pre-pubertal leukocyte depletion using sub-lethal γ-irradiation is associated with impaired ductal development and in Mϕ-deficient *Csf1*^*op*^/*Csf1*^*op*^ mice, misshapen TEBs fail to properly invade the mammary fat pad at the rate observed in age-matched controls (Gouon-Evans et al., 2000; Ingman et al., 2006; Van Nguyen and Pollard, 2002). Previous studies analyzing Mϕ density and distribution in mouse mammary tissue sections have shown recruitment of F4/80^+^ Mϕs to the pubertal epithelium and their convergence around the neck of TEBs (Gouon-Evans et al., 2000; Schwertfeger et al., 2006), where adult mammary stem/progenitor cells are thought to reside (Lloyd-Lewis et al., 2017; Sreekumar et al., 2015).

3D imaging of mammary tissue from pubertal *Csf1r-EGFP* mice revealed that mammary TEBs were enveloped by Mϕs, with spatial clustering observed (**Fig. 2A** and **Supplementary Fig. 4A**). Previous studies using the F4/80 marker indicated that Mϕs were restricted to the neck of TEBs, whereas eosinophils (distinguished by their eosinic cytoplasm and segmented nuclei) were concentrated at the TEB head (Gouon-Evans et al., 2000, 2002). By contrast, in this study GFP^+^ Mϕs in both locations shared stellate morphology (**Fig. 2A** **and Supplementary Fig. 4A**) and neither showed any evidence of segmented nuclei (**Supplementary Fig. 4A**). A small number of mammary Mϕs were observed inside the body of TEBs (**Fig. 2A**), where they may contribute to clearance of apoptotic cells from the TEB lumen (Gouon-Evans et al., 2000; Humphreys et al., 1996; Paine and Lewis, 2017). GFP^+^ Mϕs were found along the length of the ductal epithelium in the pubertal gland (**Fig. 2B** and **Supplementary Fig. 4B**) and in some cases appeared to be positioned between the luminal and basal cell layers (**Fig. 2B**, arrow). Intraepithelial Mϕs, detected with F4/80, are a feature of ductal epithelia throughout the body (Hume et al., 1984). It is currently unclear how these interposed Mϕs affect luminal-basal cell connections [e.g., desmosomes and gap junctions (Shamir and Ewald, 2015)] and their precise function within the epithelial bilayer. GFP^+^ cells were also dispersed throughout the mammary fat pad (**Fig. 2** and **Supplementary Fig. 4**) (Chua et al., 2010; Schwertfeger et al., 2006) and were densely packed in the inguinal lymph node (**Fig. 2C** and **Supplementary Fig. 4B**) and nipple region (**Fig. 2D**).

**Fig. 2:**
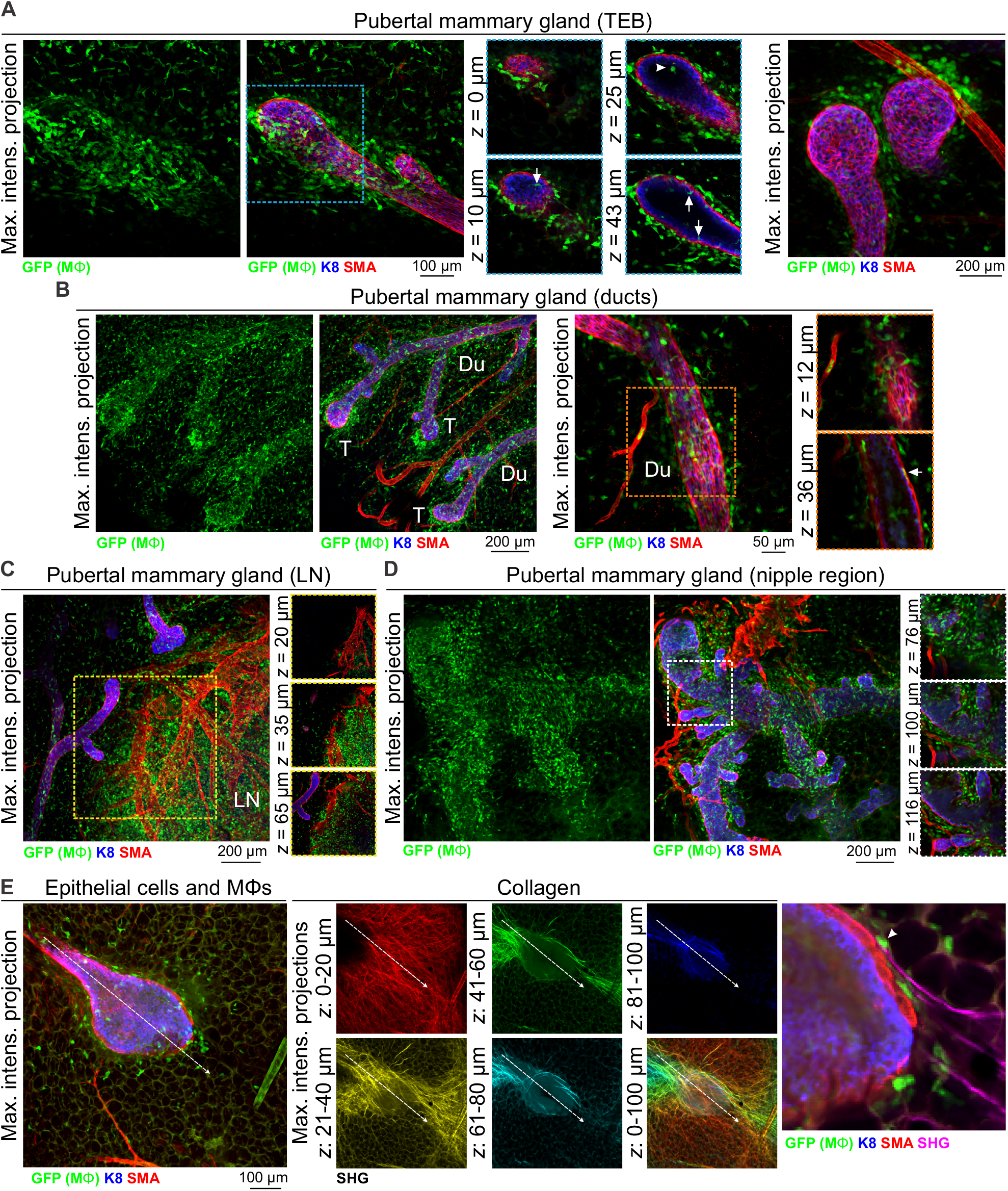
Mϕs in the mammary glands of pubertal virgin mice. Maximum intensity *z*-projection and individual optical slices of cleared mammary tissue from pubertal (6-7 week old) *Csf1r-EGFP* mice. K8 immunostaining reveals the luminal cell layer; SMA marks the basal cell layer and SMA-positive vessels. (**A**) terminal end buds (TEBs), (**B**) ductal regions, (**C**) inguinal lymph node and (**D**) nipple region. Arrows in (A) show Mϕs that have invaded the TEB epithelium and lumen (arrowhead). Arrow in (B) shows a Mϕ positioned between the epithelial bilayer. T, ductal tips; Du, ducts; LN, lymph node. Images are representative of 3 mice. (**E**) Second harmonic generation (SHG) showing fibrillar collagens around a TEB structure. Image stacks in middle panel are depth-coded (R-Y-G-C-B). Dashed arrow shows direction of TEB growth. Arrowhead in (E) shows a Mϕ interacting with collagen.

Mammary Mϕs have been shown to organize collagen into fibrillar bundles to steer TEB growth through the stromal fat pad (Ingman et al., 2006). We therefore examined fibrillar collagens with second harmonic generation (SHG) (Williams et al., 2005) in tissue from *Csf1r-EGFP* mice at depth using an immersion-based optical clearing approach, which preserves endogenous fluorescence and tissue architecture (Lloyd-Lewis et al., 2016; Vigouroux et al., 2017). Although surface collagen fibers in the mammary gland were dense and multi-directional [**Fig. 2E** (red)], deeper collagen fibers proximal to the growing TEB were aligned along its perimeter, extended in the direction of TEB growth and were associated with Mϕs (**Fig. 2E**). These data provide further evidence that mechanical forces from the stroma guide epithelial development in the normal mammary gland (Ingman et al., 2006; Stewart et al., 2019).

### 3.3 Mϕs are intimately associated with the mature ductal epithelium

Mϕs are present in the post-pubertal mouse mammary gland at all phases of the estrus cycle, with the numbers being highest in diestrus (Chua et al., 2010). In tissue sections at all estrus stages, F4/80^+^ cells are detectable around alveolar side buds versus ducts, where they are thought to promote the development and regression of these transient structures (Chua et al., 2010). Using 3D imaging of mammary tissue from *Csf1r-EGFP* mice, we observed similar numbers of Mϕs closely-associated with mammary ducts (**Fig. 3A** and **Supplementary Fig. 5**) and side buds (**Fig. 3B** and **Supplementary Fig. 5A**). As in the pubertal epithelium, Mϕs were also positioned between the luminal and basal cell layers in mature ducts and buds (**Fig. 3A-B** and **Supplementary Fig. 5B**, arrowheads) with some evidence of periodicity in intraepithelial Mϕ placement (**Supplementary Fig. 5B**). This is consistent with regular distributions of Mϕs in many locations throughout the body (Hume et al., 2019b). SHG of mature ducts revealed some fibrillar collagens that were located around the ducts and vessels (**Supplementary Fig. 5C**).

**Fig. 3:**
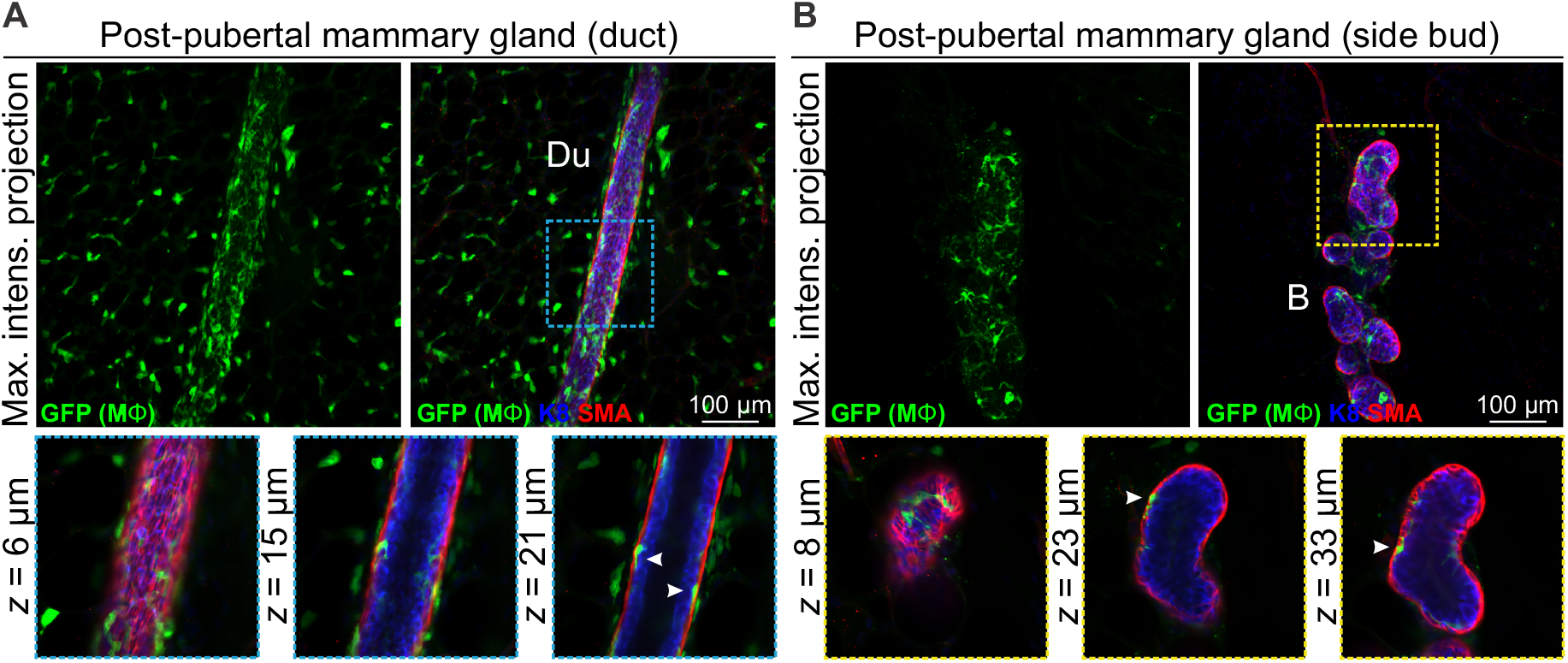
Mϕs in the mammary glands of post-pubertal virgin mice. Maximum intensity *z*-projection and individual optical slices of cleared mammary tissue from post-pubertal (12 week-old) *Csf1r-EGFP* mice. K8 immunostaining shows luminal cells; SMA immunostaining reveals basal cells and SMA-positive vessels. (**A**) Mammary ducts and (**B**) side buds. Du, duct; B, side bud. Arrowheads show Mϕs that are positioned within the epithelial bilayer. K8 immunostaining reveals the luminal cell layer and SMA marks the basal cell layer. Images are representative of 3 mice.

### 3.4 Mϕs surround alveolar units in gestation and lactation

Mϕ deficient *Csf1*^*op*^/*Csf1*^*op*^ female mice have compromised fertility (Pollard et al., 1991). Amongst those that do generate offspring, none are able to nurture a full litter, despite normal maternal behaviors (Pollard and Hennighausen, 1994). In-depth analyses of mammary tissue from pregnant and lactating *Csf1*^*op*^/*Csf1*^*op*^ mice showed incomplete branching and precocious alveolar development (Pollard and Hennighausen, 1994) and F4/80^+^ cells have been detected around the developing and functional alveolar units during pregnancy and late gestation (Gouon-Evans et al., 2002).

3D analysis of mammary tissue from pregnant *Csf1r-EGFP* mice (day 14.5 gestation, dG) confirmed Mϕ localization around the expanding alveolar structures (**Fig. 4A** and **Supplementary Fig. 6**). By lactation, Mϕs were observed immediately adjacent to alveolar basal cells, where they frequently imitated basal cell morphology (**Fig. 4B-C**, white arrowheads). Mϕs were also present within lactational alveoli (**Fig. 4C**, arrow), consistent with their enrichment in breast milk (Field, 2005).

**Fig. 4:**
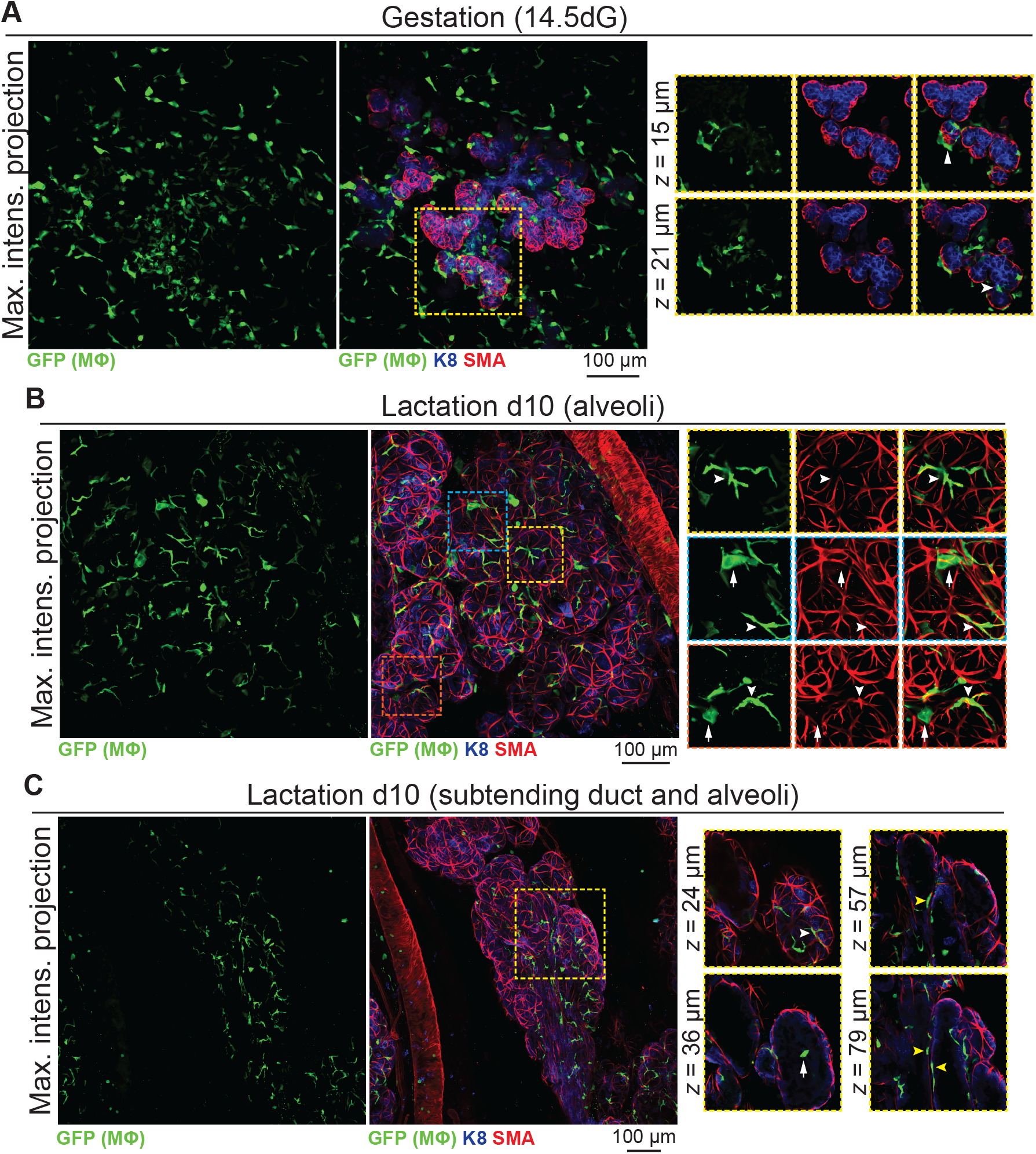
Mϕs in the mammary glands of pregnant and lactating mice. Maximum intensity *z*-projection and individual optical slices of cleared mammary tissue from (**A**) pregnant (14.5 days gestation, dG) and (**B-C**) lactating (day 10 lactation, d10) *Csf1r-EGFP* mice. K8 immunostaining reveals K8-positive luminal cells; smooth muscle actin (SMA) marks the basal/myoepithelial cells and SMA-positive vessels. Arrowheads in (A) show Mϕs that are interacting with the developing alveolar epithelium. In (B and C), white arrowheads show Mϕs that are aligned along basal cells (versus white arrows showing Mϕs that are not imitating basal cell morphology). Yellow arrowheads in (C) show Mϕs that are positioned between the ductal epithelial bilayer. Images are representative of 3 mice at each developmental stage.

### 3.5 The irreversible phase of involution is associated with an increase in Mϕnumber in and around regressing alveolar structures

The number of Mϕs surrounding the mammary epithelium increases drastically from days 3-4 of involution (Hughes et al., 2012; Lund et al., 1996; Stein et al., 2004), and involution-associated Mϕs appear polarized toward tissue repair (O’Brien et al., 2010). The recruitment and polarization of Mϕs in the involuting mammary gland is regulated by epithelial *Stat3* expression (Hughes et al., 2012). Moreover, pre-weaning depletion of CSF1R-expressing cells reduces mammary epithelial cell death during post-lactational involution, an effect that can be reversed by orthotopic transplantation of bone marrow-derived Mϕs (O’Brien et al., 2012).

To further examine Mϕ number, morphology and distribution in the regressing mammary gland in 3-dimensions, we analyzed optically-clear tissue from *Csf1r-EGFP* mice during the irreversible phase of involution. Relative to other developmental stages, Mϕ density was high at 96 h involution and Mϕs were observed around and inside ducts and regressing alveoli (**Fig. 5A-B**). Large aggregates of GFP^+^ cells, reminiscent of homotypic fusion (MacLauchlan et al., 2009), were also observed inside degenerating alveolar structures (**Fig. 5B** arrowheads). Similar aggregates of GFP^+^ Mϕs have been observed in a model of epithelial regeneration in the kidney following transient ischemia (Joo et al., 2016).

**Fig. 5:**
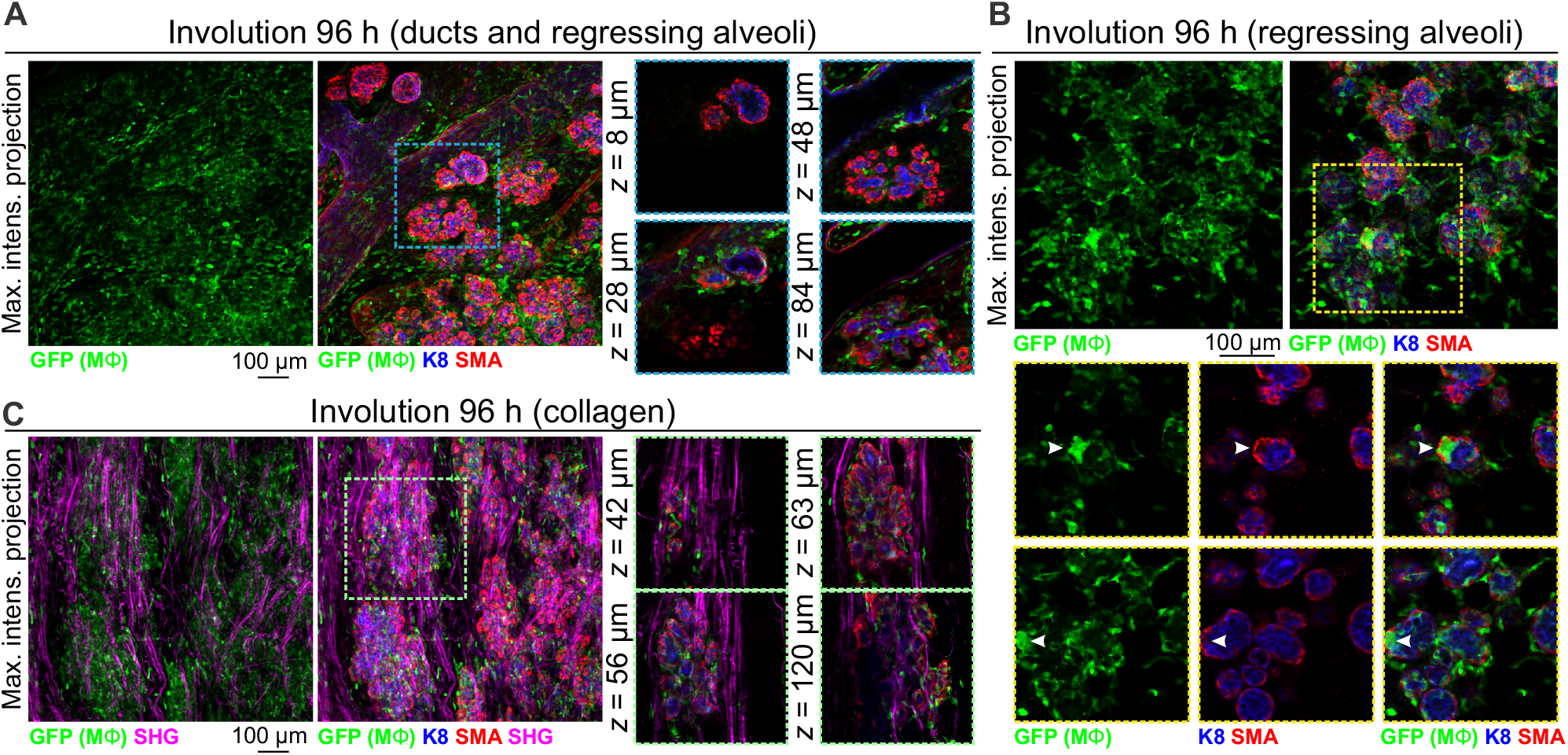
Mϕs in the mammary glands of mice during post-lactational involution. (**A-C**) Maximum intensity *z*-projection and individual optical slices of cleared mammary tissue from *Csf1r-EGFP* mice during involution (96 h post forced weaning). K8 immunostaining shows luminal cells; SMA immunostaining reveals basal cells and SMA-positive vessels. Arrowheads in (B) show a cluster of GFP^+^ Mϕs inside of collapsed alveolar units. (**C**) SHG showing fibrillar collagens surrounding regressing alveoli. Images are representative of 3 mice.

Collagen density increases during mammary gland involution and partially-degraded nonfibrillar collagens have been suggested to be chemotactic for Mϕs (O’Brien et al., 2010). Intra- and interlobular fibrillar collagens were observed with SHG in *Csf1r-EGFP* mice and GFP^+^ Mϕs were observed to be associated with collagen fibrils (**Fig. 5C**).

## 4 Discussion

Mϕs contribute to mammary gland development and remodeling at all developmental stages (Chua et al., 2010; Dai et al., 2002; Gouon-Evans et al., 2000; Hughes et al., 2012; Ingman et al., 2006; O’Brien et al., 2010, 2012; Pollard and Hennighausen, 1994; Van Nguyen and Pollard, 2002). In this study, we provide new insights into the relative number, morphology and distribution of Mϕs in the embryonic, pre-pubertal, pubertal, post-pubertal, pregnant, lactating and involuting mammary glands of fluorescent reporter-positive mice *in situ* in 3-dimensions (**Fig. 6**). Our study yields a number of important observations that could only be revealed by multi-dimensional imaging using a tamoxifen-independent, cell type-specific fluorescent reporter model (Hume et al., 2019b, 2019a). Firstly, in contrast to previous reports (Gouon-Evans et al., 2000, 2002), we demonstrate that Mϕs encase the length of elongating TEBs and are not restricted to the TEB neck. Mammary Mϕs were also frequently embedded between luminal and basal cells of the ductal epithelium. This has previously been observed in mammalian ductal epithelia, including the bile duct, salivary gland, tracheobronchial gland and mammary gland using thin sections prepared from formalin-fixed paraffin-embedded or frozen tissue (Hume et al., 1984; Sun et al., 2013). Regularity in the spacing of these intraepithelial Mϕs was also noted. Our findings suggest a close functional relationship between Mϕs and ductal epithelial cells, and possible communication between morphologically-related Mϕ populations. Further studies are needed to determine whether these intraepithelial Mϕs share similar gene and protein expression patterns and whether this information can be used to probe their function, retention and passage within the epithelium. Tissue Mϕs have been shown to be influenced by properties of their specific niche within each tissue (e.g., anchoring scaffolds and local cues) (Chakarov et al., 2019; Mondor et al., 2019). Single cell sequencing of isolated mammary Mϕs from *Csf1r-EGFP* mice at distinct developmental stages, as exemplified by recent studies of other tissues (Chakarov et al., 2019; Mondor et al., 2019), might help to reveal the extent of functional diversity within Mϕ populations in this organ. Secondly, we show that Mϕs alter their morphology at distinct developmental stages, including the transition from gestation to lactation. The localization of Mϕs around growing alveolar units during gestation and the observation that Mϕ-deficient *Csf1*^*op*^/*Csf1*^*op*^ mice exhibit precocious alveolar development, suggests that during this phase, alveolar-associated Mϕs may restrain alveologenesis. During lactation, Mϕs altered their anatomical position and were observed to closely imitate the morphology of adjacent, differentiated alveolar basal cells. Whether these cells specifically align themselves with oxytocin-responsive basal cells during lactation to modify basal cell function (Davis et al., 2015; Stevenson et al., 2019) or more simply to occupy the physical space that these force-exerting cells create within the alveolar epithelium (Davis, 2016; Stewart et al., 2019), remains unknown. Such a function might be analogous to the role of CSF1-dependent Mϕs in the regulation of peristalsis in the muscularis externa of the intestine (Muller et al., 2014).

**Fig. 6:**
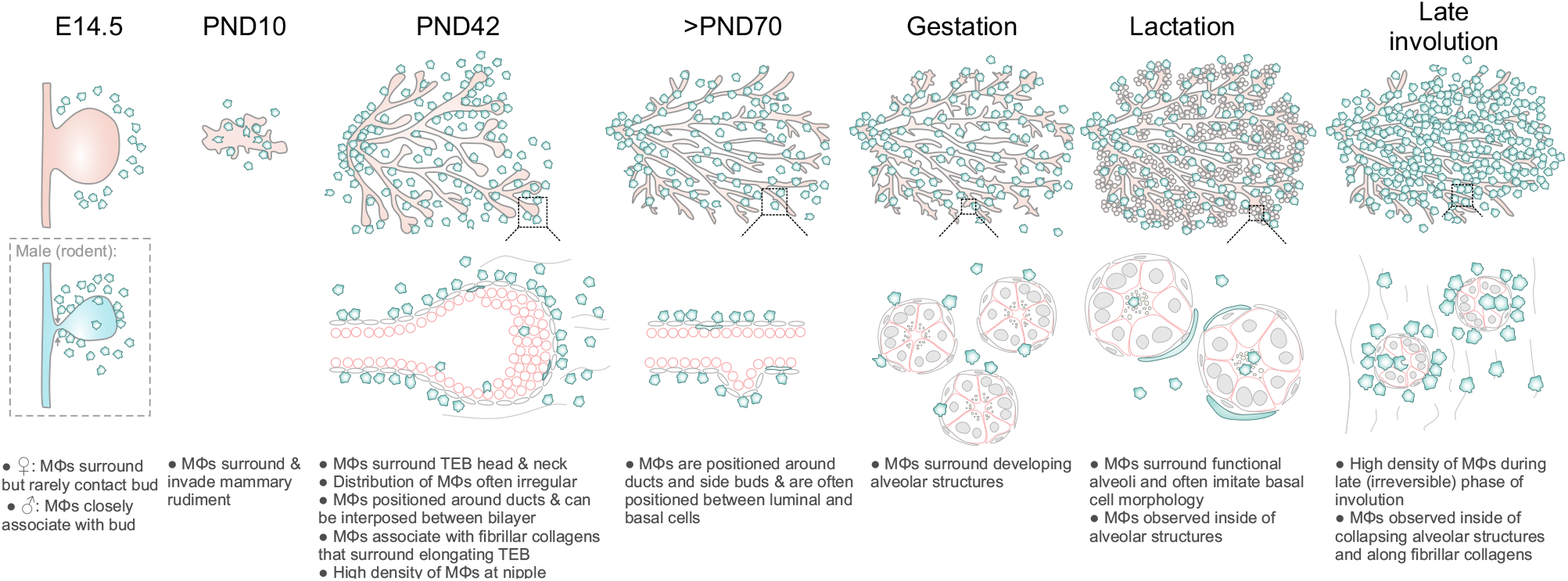
Diagram summarizing Mϕ distribution in the mouse mammary gland during distinct phases of development and remodeling.

Finally, we were able to visualize for the first time tissue-resident Mϕs in the mesenchyme surrounding the mammary epithelial bud in 14.5 day-old female embryos. Intriguingly, these embryonic Mϕs rarely contacted the epithelial cells of the developing mammary bud at this stage of embryogenesis. This is in striking contrast to epithelial-Mϕ interactions in the early postnatal period, where Mϕs surround and invade the rudimentary ductal epithelium. This also contrasts with the male embryo, where Mϕs were often observed to both contact and infiltrate the epithelial bud at the time when its connection to the overlying epidermis is severed and the structure begins to regress (Cowin and Wysolmerski, 2010; Dunbar et al., 1999; Heuberger et al., 2006). At this stage, Mϕs may have an important role in clearing apoptotic epithelial and mesenchymal cells (Dunbar et al., 1999; Henson and Hume, 2006).

Mammary stem/progenitor cells are located within the mammary bud (in the embryo) and TEBs (in puberty). After ductal elongation is complete and TEBs regress, however, the location of long-lived mammary stem/progenitor cells and their putative niche remains unknown, although it has been suggested that these cells are deposited along the ductal epithelium by elongating TEBs (Davis et al., 2016; Lloyd-Lewis et al., 2017). In the embryo, Mϕs were positioned uniformly around, but not in contact with, the mammary bud. These data suggest that if a mammary stem cell-macrophage niche exists in the embryo around the time of lineage segregation, it operates over the scale of tens of micrometers and is fairly homogeneous. Mϕs were also positioned around pubertal TEBs, however, in contrast to the embryo, these cells contacted and infiltrated TEBs, were more densely arranged around these structures and often showed spatial clustering. Future studies combining tamoxifen-independent *Dll1-mCherry* (Chakrabarti et al., 2018) and *Csf1r-EGFP* mouse models with optical tissue clearing and 3D imaging may help to reveal the precise location of mammary stem/progenitor cells within TEBs and the post-pubertal ductal epithelium. It should be noted, however, that whilst ductal elongation is delayed in *Csf1*^*op*^/*Csf1*^*op*^ mice, these structures are still capable of invading the fat pad and by 12 weeks of age have reached the fat pad limits (Gouon-Evans et al., 2000). These findings imply that mammary epithelial cells have mechanisms to overcome insufficiencies in niche signaling. One candidate is the alternative CSF1R ligand, IL34, which may also be expressed by mammary epithelial cells (DeNardo et al., 2011). Studies investigating the activation and roles of the CSF1R in mammary development have been thwarted by the severe postnatal phenotype of *Csf1r*^−^/*Csf1r*^−^mice (Chitu and Stanley, 2017), but may be more amenable to study in recently described *Csf1r*^−^/*Csf1r*^−^ rats (Pridans et al., 2019). Alternatively, these findings may reflect a long-term plasticity in mammary epithelial cells (Lilja et al., 2018) and a shifting definition of ‘stemness’ in some tissues away from a unidirectional, top-down model to a model where stemness is considered as a cell state that may be acquired or extinguished under specific microenvironmental conditions (Laplane and Solary, 2019). A closer examination of mammary cell behaviors—including lineage segregation—under conditions of Mϕ depletion may provide important insights into epithelial plasticity in this vital mammalian organ.

## 6 Author Contributions

F.M.D. and T.A.S. performed all experiments. F.M.D., D.A.H. and T.A.S. conceived and designed the experiments. K.H., T.A.S., D.A.H. and F.M.D. analyzed the results. F.M.D. wrote the manuscript. D.A.H., K.H. and T.A.S. edited the manuscript.

## 8 Acknowledgments

The authors acknowledge the Translational Research Institute (TRI) for the research space, equipment and core facilities that enabled this research. We thank the UQ Biological Resource staff for animal care and husbandry; Mr Alex Stevenson for laboratory management and ordering of consumables; A/Prof Allison Pettit and Dr. Katharine Irvine for their helpful comments on the manuscript; Dr. Jerome Boulanger (MRC Laboratory of Molecular Biology) for the 3D de-noising algorithm; and Mr Eric Pizzani (Translational Research Institute) for research computing support.

