## Supplementary figures for "Developmental stage-specific distribution of macrophages in mouse mammary gland"

Supplementary Material

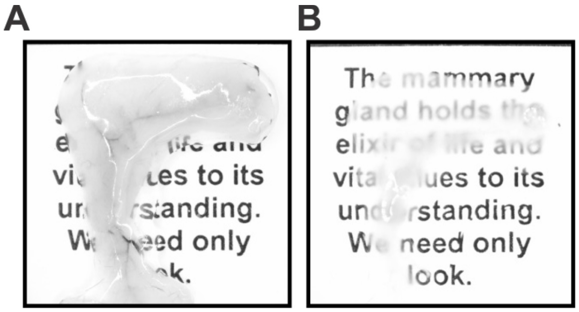


**Supplementary Figure 1: Optical tissue clearing in the mammary gland using a modified CUBIC protocol.** Images show before (**A**) and after (**B**) clearing.

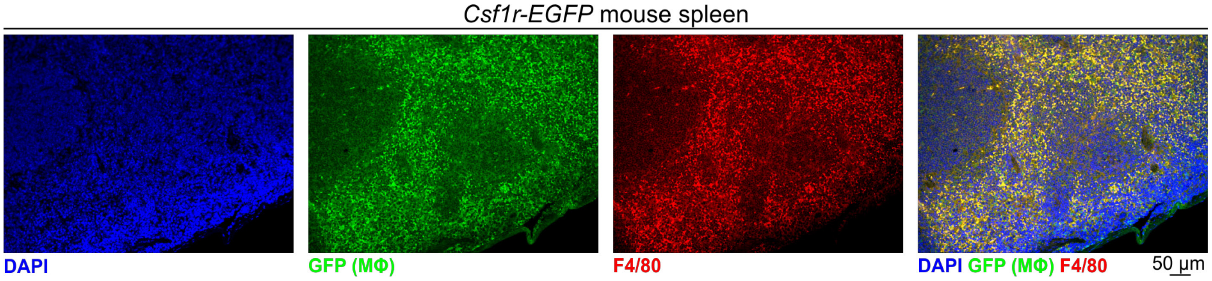


**Supplementary Figure 2: Analysis of GFP$^{+}$ cells in the spleen.** Tissue section from *Csf1r-EGFP* mouse immunostained with F4/80 and GFP antibodies.

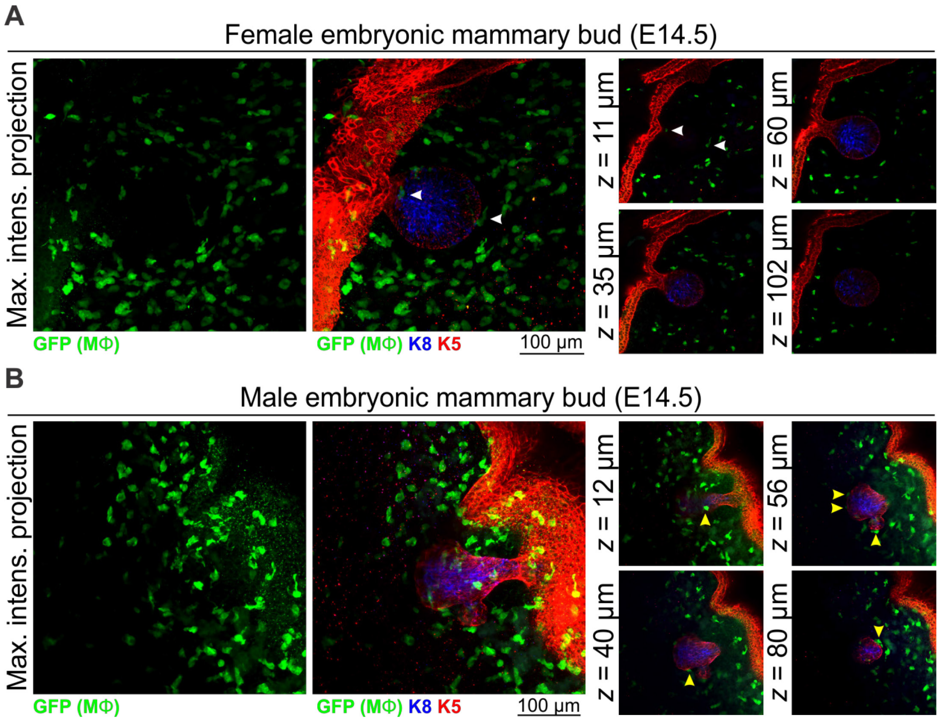


**Supplementary Figure 3: Additional images of embryonic mammary Mϕs.** Maximum intensity *z*-projection and individual optical slices of cleared tissue from (**A**) E14.5 female embryos and (**B**) E14.5 male embryos. White arrowheads in (A) point to Mϕs that appears to be in contact with the embryonic bud in the maximum intensity projection, but are revealed to be positioned in the mammary mesenchyme around the bud in optical slices. Yellow arrowheads in (B) point to Mϕs that are in direct contact with the embryonic bud. Related to Figure 1.

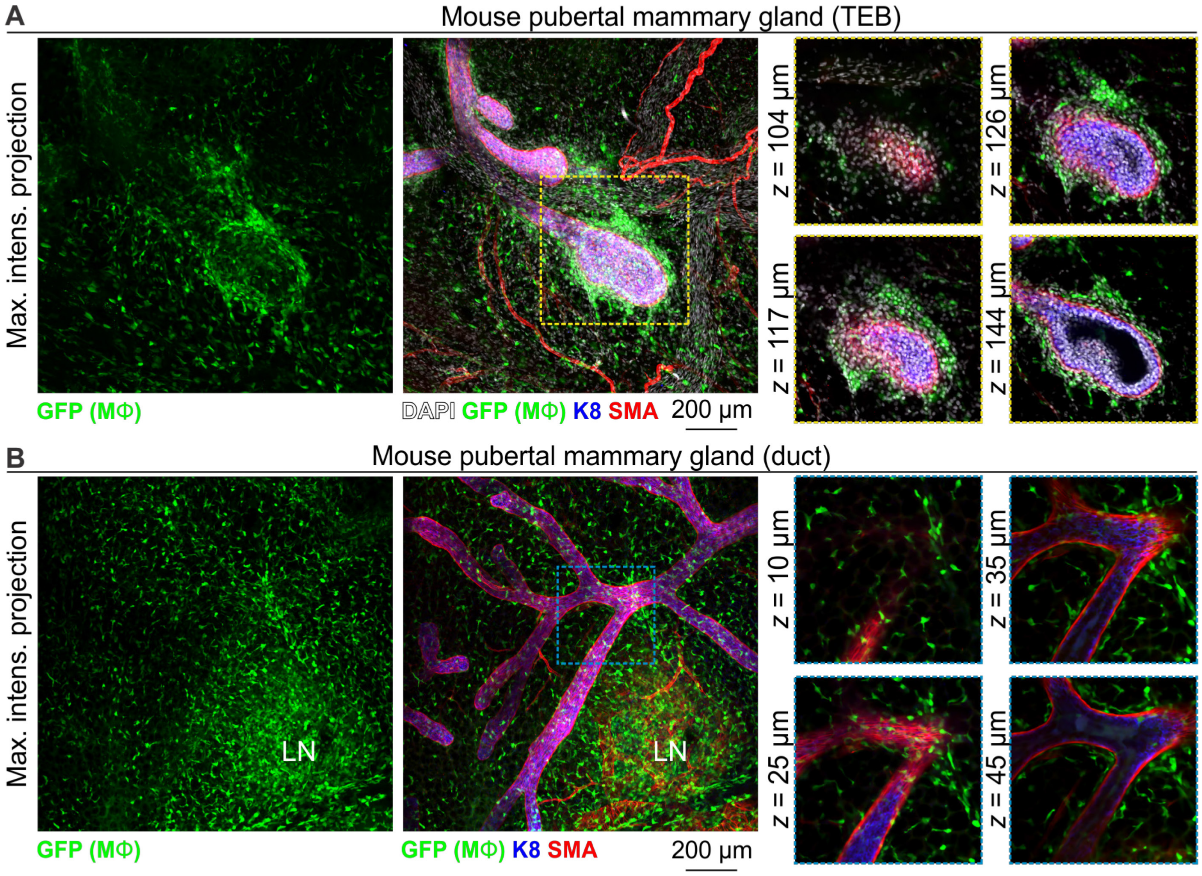


**Supplementary Figure 4: Additional images of Mϕs in the mammary glands of pubertal virgin mice.** Maximum intensity *z*-projection and individual optical slices of cleared mammary tissue from pubertal (6-7 week old) *Csf1r-EGFP* mice. (**A**) Terminal end buds (TEBs) and (**B**) ductal regions. LN, lymph node. Related to Figure 2.

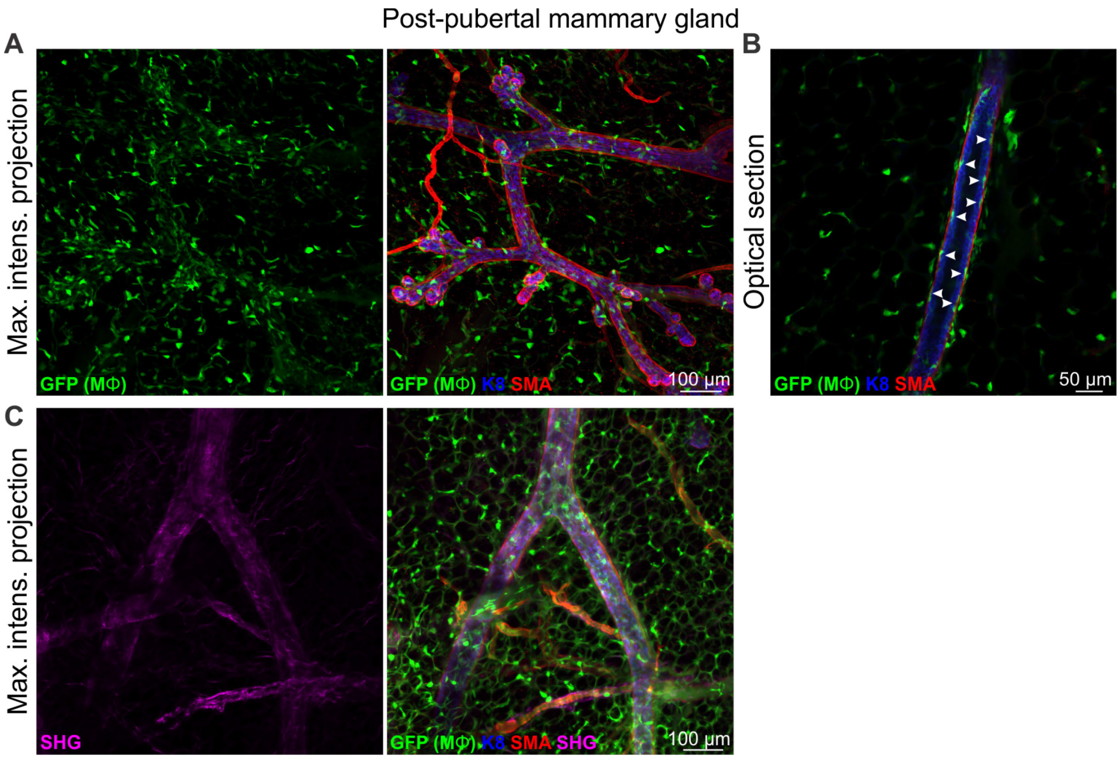


**Supplementary Figure 5: Additional images of Mϕs in the mammary glands of post-pubertal virgin mice.** (**A-C**) Maximum intensity *z*-projection and individual optical slices of cleared mammary tissue from post-pubertal (12 week-old) *Csf1r-EGFP* mice. Arrowheads in (B) show periodic spacing of intraepithelial Mϕs (duct shown in Fig. 3). Related to Figure 3.

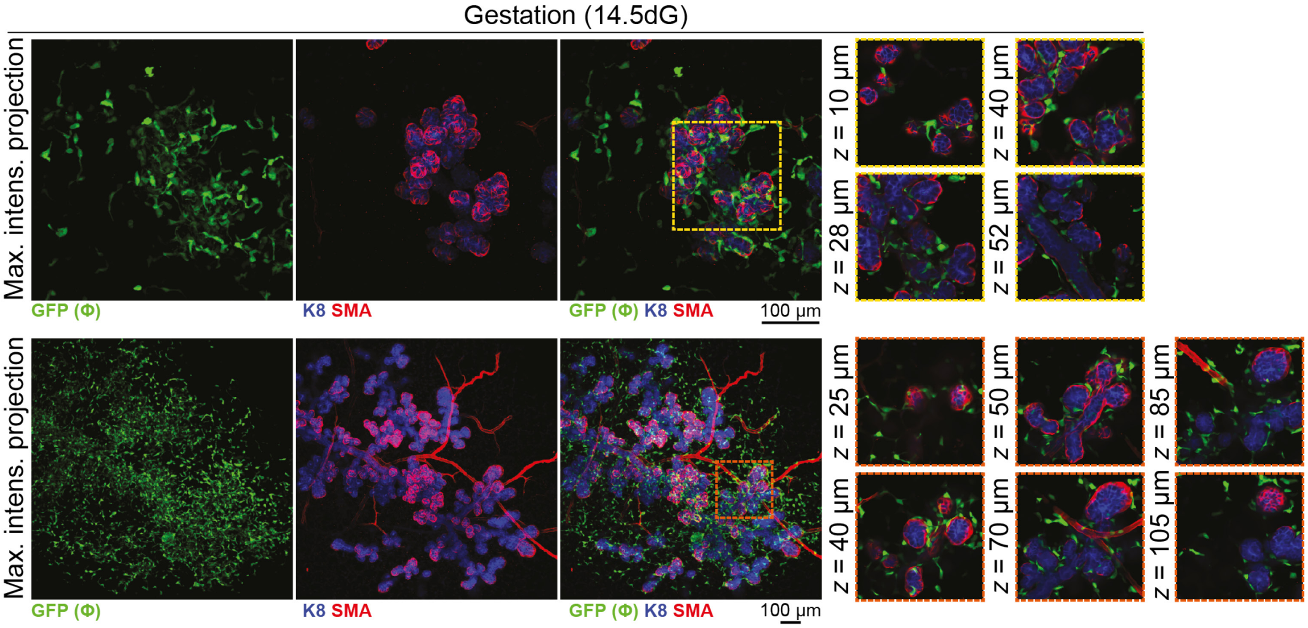


**Supplementary Figure 6: Additional images of Mϕs in the mammary glands of pregnant mice.** Maximum intensity *z*-projection and individual optical slices of cleared mammary tissue from pregnant *Csf1r-EGFP* mice. Related to Figure 4.
